# Modeling Reactive Species Metabolism in Colorectal Cancer for Identifying Metabolic Targets and Devising Therapeutics

**DOI:** 10.1101/2022.05.03.490417

**Authors:** Prerna Bhalla, Subasree Sridhar, Justin Kullu, Sriya Veerapaneni, Swagatika Sahoo, Nirav Bhatt, GK Suraishkumar

**Affiliations:** Department of Chemical Engineering, Indian Institute of Technology Madras, Chennai - 600036, India; Department of Biotechnology, Bhupat and Jyoti Mehta School of Biosciences, Building - 1, Indian Institute of Technology Madras, Chennai - 600 036, India; International Research Initiative, Global Engagement, Indian Institute of Technology - Madras, Chennai - 600 036, India

## Abstract

Reactive species (RS) are known to play significant roles in cancer development as well as in treating or managing cancer. On the other hand, genome scale metabolic models are being used to understand cell metabolism in disease contexts including cancer, and also in planning strategies to handle diseases. Despite their crucial roles in cancers, the reactive species have not been adequately modeled in the genome scale metabolic models (GSMMs) when probing disease models for their metabolism or detection of drug targets. In this work, we have developed a module of reactive species reactions, which is scalable - it can be integrated with any human metabolic model as it is, or with any metabolic model with fine-tuning. When integrated with a cancer (colorectal cancer in this case) metabolic model, the RS module highlighted the deregulation occurring in important CRC pathways such as fatty acid metabolism, cholesterol metabolism, arachidonic acid and eicosanoid metabolism. We show that the RS module helps in better deciphering crucial metabolic targets for devising better therapeutics such as FDFT1, FADS2 and GUK1 by taking into account the effects mediated by reactive species during colorectal cancer progression. The results from this reactive species integrated CRC metabolic model reinforces ferroptosis as a potential target for colorectal cancer therapy.

## 1 Introduction

Reactive species (RS) are understood as a class of highly reactive compounds that can be generated via enzymatic/non-enzymatic reactions, predominantly comprising of reactive oxygen species (ROS), reactive nitrogen species (RNS) and reactive sulphur species (RSS) [1, 2]. These molecular entities are known to participate in various regulatory, signaling, metabolic, immune and inflammatory processes, and hence contribute to maintain cellular redox homeostasis [3, 4]. Accumulation of RS due to a mismatch between their production and degradation rates results in redox imbalance and oxidative stress.

Oxidative stress is a key characteristic of various diseases such as cancer, Alzheimer’s, and cardiovascular disorders [5]. For example, several studies have reported the involvement of RS and consequent oxidative stress in the deregulation of various metabolic and signaling pathways that result in colorectal carcinogenesis [6, 7]. Two of the most important hallmarks of cancer are altered stress response and metabolic re-wiring [8]. Oxidative stress occurs due to many conditions such as hypoxia, nutrient scarcity, etc., and the altered stress response includes deregulated signaling, DNA damage, and others [9]. Under these circumstances, metabolic re-wiring provides a selective advantage during initiation and progression of tumors to address say, scarcity in nutrient or oxygen [10]. Thus, altered stress response and metabolic re-wiring together favour survival of the tumour cells. Modelling cancer metabolism using the omics data will help in understanding metabolic re-wiring or reprogramming in the tumour [11, 12]. Constraint-based modeling (CBM) and context-specific metabolic models have found substantial applications in understanding the underlying complexity of cancer metabolism. Considering the ever-evolving metastatic nature and inter- and intra-tumour heterogeneity of the disease, these models have been extensively used to analyze cancer phenotypes, as well as to uncover potential therapeutic targets [13, 14]. Further, oxidative stress is significantly associated with premature mortality in CRC patients [15]. Given the significance of RS homeostasis in cancer initiation and progression, it would be insightful to model RS metabolism in disorders such as colorectal cancer.

In this work, to model RS metabolism, we have formulated for the first time, a RS module, consisting of metabolic reactions of various RS (ROS, RNS, RSS, and the less prevalent reactive halide species, RHS) with important biomolecules such as nucleic acids, amino acids, etc. This RS module is scalable, i.e. it can be integrated with any human metabolic model, or with some fine-tuning to any metabolic model, to investigate RS effects in a variety of situations of interest such as diseases, metabolite production, or others. Our earlier approaches to mathematically represent the effects of RS on cells through dynamic equations based on fundamental kinetic expressions were severely limited by the non-availability of the relevant kinetic constants. Also, earlier approaches in the literature to model RS effects were specific say, to processes that participate in bacterial iron homeostasis [16] or plant C1 metabolism [17], whereas the approach in this work is highly scalable and applicable to any human metabolic model, or to any metabolic model with fine-tuning.

Further, the formulated RS module was integrated with the context-specific CRC model to model RS metabolism and explore the latter’s influence on cancer cell metabolism. The modeling predictions were thoroughly examined by comparing them with the existing literature findings. Furthermore, the altered metabolism of CRC upon incorporation of RS was investigated to predict metabolic therapeutic targets. Cancer cells evolve and develop certain mechanisms that alleviate the oxidative stress [18] and redox therapies against cancer can exploit this phenomenon.

## 2 Methods

### 2.1 Formulation of the reactive species module

The NDRL/NIST solution kinetics database https://kinetics.nist.gov/solution/ was a source of the RS associated reactions along with various other sources from the literature. Most of the NIST database reactions utilizing RS as reactants or generating them as products were incomplete, and hence were curated using suitable information from the literature. It was ensured that the reactions that were taken from these databases took place at physiological pH (6.8 - 7.5). The pubchem website, https://pubchem.ncbi.nlm.nih.gov/ was used to find the neutral formulae of the metabolites, and then the charges of the RS associated metabolites were obtained from https://chemaxon.com/products/marvin. The Virtual Metabolic Human database, https://www.vmh.life/ was referred to find the abbreviations of the metabolites associated with RS reactions so that they are consistent with the ones used in the Recon3D model. The resulting reactions and metabolites associated with RS were compiled and translated into the COBRA model format using the rBioNet [19] tool of the COBRA Toolbox [20]. rBioNet is an extension of the COBRA toolbox used to create metabolic reconstruction networks. Names, chemical formulae of the metabolites, charges of the metabolites, names of the reactions and balanced formulae of the reactions were all updated in the RS module. A schematic of the work done toward building and integration of RS module into the CRC and colon metabolic models is presented in Figure 1.

**Fig 1.**
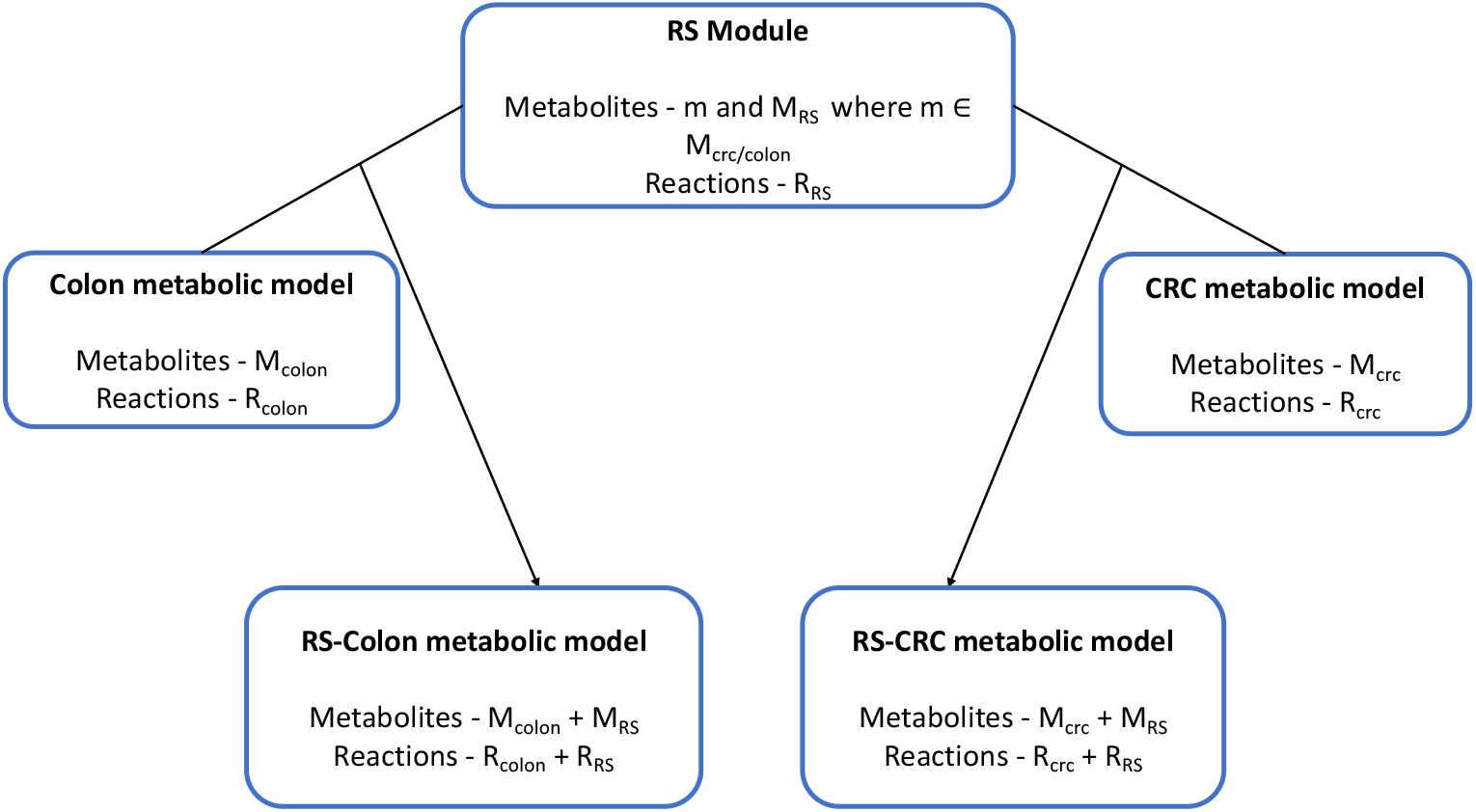
Integration of a context specific metabolic models built from Recon 3D Model and the RS module. RS module contains exclusive metabolites and reactions M_*RS*_ and R_*RS*_. Few of the metabolites, m are common to the RS module and the 2 metabolic models

### 2.2 Computational analysis

#### 2.2.1 Flux variability analysis (FVA) and flux span ratio (FSr)

The formulated RS module was integrated with DMEM-constrained context-specific models (healthy colon and CRC) [21] and the RS reactions were tested for their flux activity. The blocked (flux inactive) reactions were debugged, and the integrated models were then subjected to various analyses such as flux balance analysis (FBA), flux variability analysis (FVA), flux span ratio (FSr) analysis [21] and flux sampling to investigate the transformed metabolism of CRC in presence of RS. FBA gives the optimal flux under a given set of constraints and an objective function but there are numerous optimal fluxes for an underdetermined system like this [22]. FVA is used to find the minimum and maximum flux for reactions (range of flux) in the metabolic model with or without emphasis on an objective function [23]. FSr values are defined as the ratios of flux ranges of reactions in diseased vs. normal condition, i.e. colon cancer vs. healthy colon. For the model comparisons of RS-CRC vs. CRC and RS-CRC vs. RS-Colon, reactions with FSr values lying above 2 or below 0.8 were deemed significantly deregulated [21]. In both the cases, upregulated reactions were those with FSr values in the range [2,4], and reactions with FSr values in the interval [0.1,0.8] were classified as downregulated reactions. Besides this, the FSr results were analyzed to map the FDA approved drugs, as well as potential drug targets using information from druggable proteome from Human Protein Atlas (HPA), https://www.proteinatlas.org/humanproteome/tissue/druggable and the FSr results were also mapped onto Cancer Dependency Map https://depmap.org/rnai/index to check for their potential as suitable gene targets.

#### 2.2.2 Sampling

In addition to the FVA analysis, flux sampling was performed to understand the feasible flux solution space of a metabolic model by generating probability distributions of steady-state reaction fluxes under given constraints [24]. Unlike FBA, both FVA and sampling can be used without assuming a particular cellular objective. One major advantage of sampling over FVA is that it gives the feasible flux values. Artificial Coordinate Hit and Run (ACHR) was used to sample fluxes from the four models, CRC, RS-CRC, colon and RS-Colon. One thousand sample points for each reaction were drawn from the four models. Flux sampling [25] estimation was done using Cobrapy [26] using the cplex solver. Two-sample Kolmogorov Smirnov Test (KS Test) in MATLAB was used to differentiate the two flux samples of each reaction while comparing CRC vs. RS-CRC and RS-Colon vs. RS-CRC and CRC vs. Colon models as well [27]. A significance level threshold of 0.05 was used for filtering the deregulated reactions in the three cases. All the deregulated reactions from FSr analysis were found to be significantly different based on the KS test and these reactions were used for further analysis.

#### 2.2.3 Production rates of metabolites

In a steady state genome scale model, there is no accumulation of metabolites because the production rates of metabolites are assumed equal to their consumption rates. Therefore, production rates of extracellular metabolites were mathematically calculated from the mean of sampled fluxes and their stoichiometric coefficients in the reactions in which they participate. Ten thousand (10000) sample points were drawn for each reaction of the 4 models to calculate the mean of the sampled fluxes [25]. The production rates of extracellular metabolites were compared between RS-CRC and RS-Colon. The significant metabolites with log fold change of *≥* 1.5 or *≤* −0.65 were considered to be upregulated and downregulated respectively [28] in the RS-CRC model. The results were compared with various serum metabolomic studies on colorectal cancer to validate the findings [29, 30].

#### 2.2.4 Lethality and synthetic lethality analysis for finding drug targets in CRC

Lethality analysis on genes and reactions to compute the single and double gene pairs and single and double reaction pairs, the deletion of which result in the reduction in the biomass, was carried out using fastfl [31] algorithm. This will be useful in predicting suitable drug targets for CRC. The algorithm takes into account the gene protein reaction (GPR) rules of the COBRA metabolic models to determine gene targets. The biomass reaction is a demand reaction without any GPR association whereas the precursors of the biomass reactions are produced and consumed through a complex network of reactions with GPR rules associated with them. The association between genes for a reaction is given in the form of OR or AND rules depending on whether the genes are the subunits of an enzyme or whether they are the isozymes of enzymes or act synergistically in regulating a reaction. This attribute can be probed to identify probable gene targets. Double gene/reaction pairs are those where the individual gene/reaction in the pair is non-essential to the biomass growth, but the gene/reaction when acting in pairs will be essential for the cell to grow, and deletion of the pair induces biomass reduction. There are many reactions in the Recon 3D models that are not provided with any gene information and such reactions are called orphan reactions [20]. Thus it becomes important to study the effects of reaction deletions in addition to the gene deletion studies. Single and double gene and reaction deletion results are provided in the supplementary file.

#### 2.2.5 DEMETER: A cancer dependency map framework

The in-silico gene drug targets for CRC as obtained from gene deletion and FSr analysis were further validated using cancer dependency map (DEMETER) https://depmap.org/rnai/index. The DEMETER database [32] registers the experimental gene knockdown effects on various cancer cell lines. This database contains quantitative data of log-fold change in cell number due to gene knockdown based on three large-scale RNAi screening datasets, namely the Broad Institute Project Achilles, Novartis Project DRIVE, and the Marcotte et al. breast cell line dataset across 712 cancer cell lines. A gene is crucial to the colon cancer growth if the log fold change value is negative [32]. Potential gene targets obtained from our analyses were validated using DEMETER log fold change values for the CRC cell line named HUTU-80.

In summary, the computational analyses performed on the RS integrated models are shown in Figure 2.

**Fig 2.**
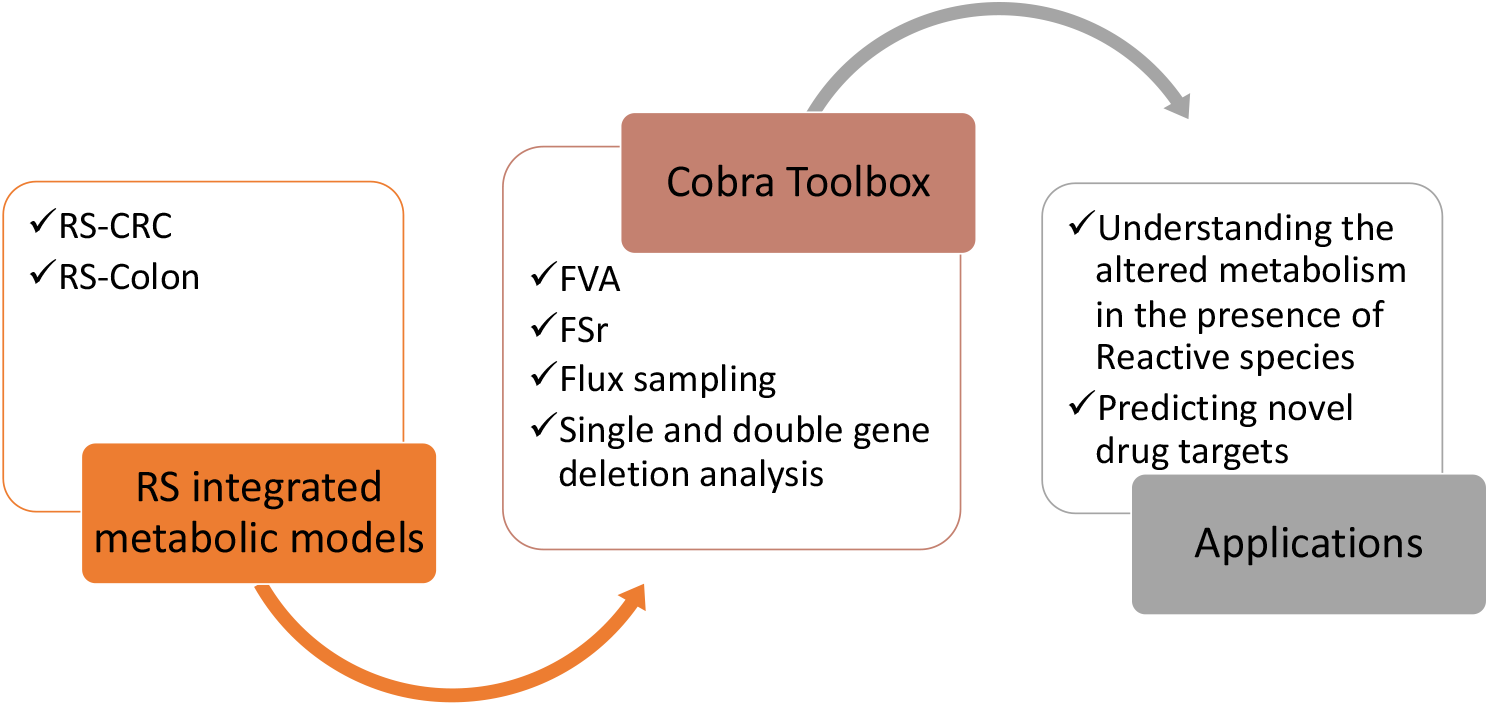
COBRA Toolbox used to perform mathematical analyses such as FVA, FSr, etc., to understand the contribution of RS in altering disease metabolism as well as in predicting drug targets

## 3 Results

### 3.1 Formulating the RS reactions paradigm as a part of the human metabolism

The existing human metabolic reconstructions such as Recon 3D [33] and HUMAN-GEM [34] do not address the effects of reactive species, except for a few mitochondrial and cytosolic ROS detoxification reactions. Therefore, the prime objective of the present work was to assemble a reactive species (RS) module pertaining to human metabolism, based on information available in literature and databases. The module was constructed by including almost all the physiologically relevant reactive species reactions of the human body under physiological conditions (details in the methods section). The current version of this module consists of 397 reactions which includes metabolic reactions (135), demands (1), exchanges (65), and transports (129) that are essential for the completeness of RS metabolism. The breakdown of the metabolic reactions as per the type of reactive species involved (ROS, RNS, RSS, etc.) are given in Table 5. Moreover, an additional 62 reactions from the Recon 3D reconstruction which are missing in the curated Recon 3D model, were included in the integrated models to enable the flow of metabolites between the CRC and colon model reactions and the RS module reactions https://www.vmh.life/. The module was built as a COBRA model with 397 reactions and 458 metabolites.

The RS module can be integrated with any metabolic model using the mergeTwoModels code available in COBRA Toolbox documentation [20]. The RS integrated metabolic models can be used to analyse the impact of RS species on any metabolic system (Figure 3). The integration of this module with CRC metabolic model has captured the in-vivo metabolic deregulation occurring in CRC better than the CRC model without the RS module integrated to it and the results are discussed in section 3.2. Further analyses on the models were carried out to understand the metabolic deregulation and to identify metabolic targets.

**Fig 3.**
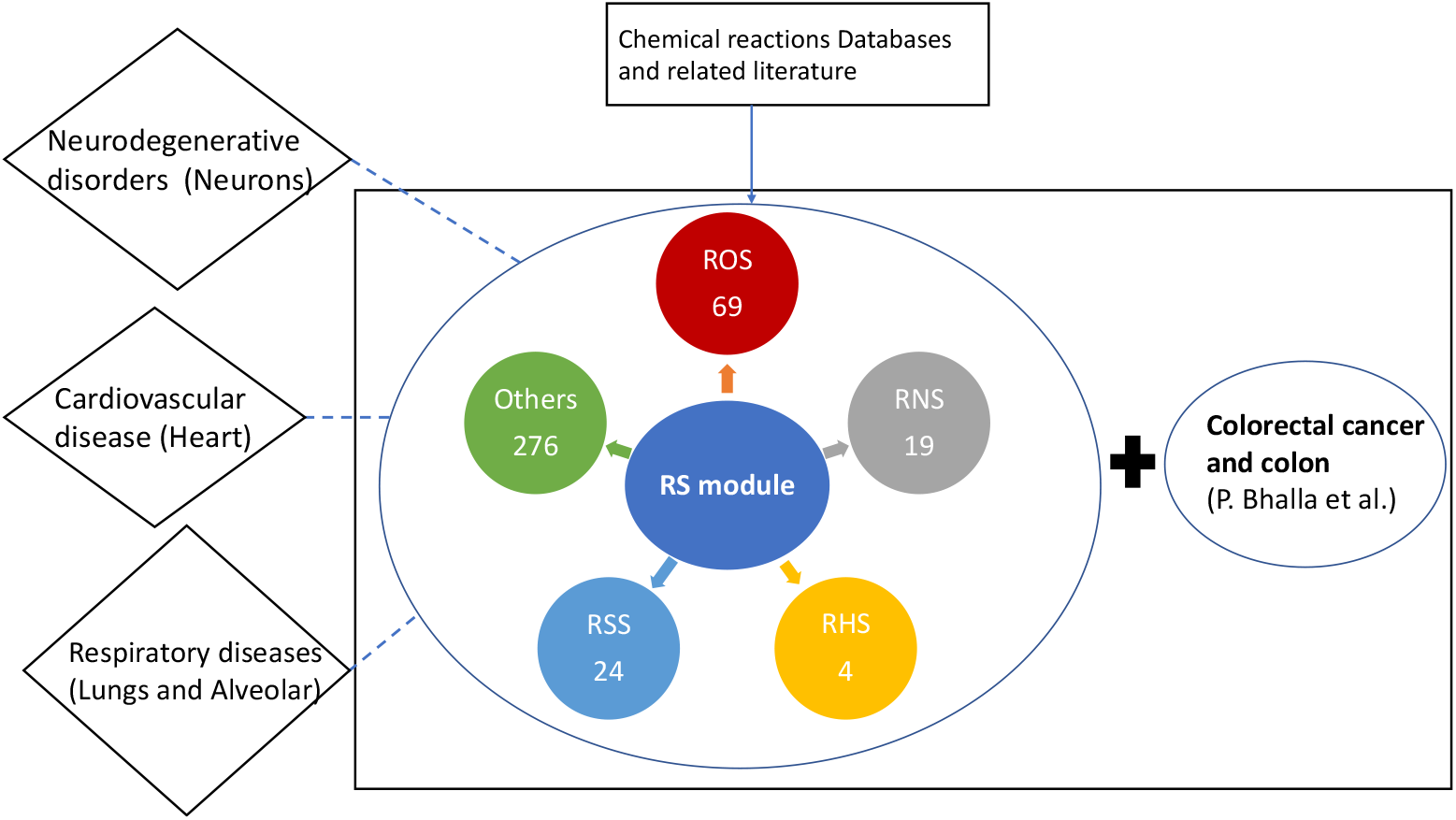
A schematic showing the integration of RS module with metabolic models of various diseases. The compiled RS module can be integrated with heart, lung, or neuronal metabolic models to study its relevance in diseases such as COVID-19, Alzheimer’s, and others. In the present study, RS module was integrated with CRC metabolic model with the aim of identifying novel drug targets.

### 3.2 Prediction of diagnostic biomarkers and glutathione redox status in the RS-CRC model

Acetate, glycine, lactate, phenylalanine, cholesterol, 3-hydroxybutyrate, histidine, isoleucine, leucine, serine, sphinganine and glucose were found to be upregulated in RS-CRC model compared to RS-Colon model and metabolites such as citrate, creatinine and glycocholic acid were found to be downregulated [29, 30]. Novel findings from this analysis on RS-CRC model are some of the upregulated metabolites that belong to RS metabolism like 4-chlorocatechol OH-adduct, guanine OH-adduct, 6-methyluracil-5-OH-adduct, purine OH-adduct, radicals from arabinose, nitrous acid and nitrogen trioxide. Some of the downregulated RS module related metabolites are sulfhydryl dimer radical anion, sulfiyl radical, nitrogen dioxide radical, carbondioxide radical anion, 2-deoxycytidine OH-adduct, 2-(methylthio)ethanol OH-adduct and 2-(methylthio)ethanol. These upregulated and downregulated RS metabolism related metabolites can serve as clinical biomarkers.

**Table 1.** Model statistics for the RS module.

| S.No | Reaction category | Number of reactions |
| --- | --- | --- |
| 1 | Metabolic RS reactions | 135 |
| 1.1 | Reactive oxygen species | 70 |
| 1.2 | Reactive nitrogen species | 23 |
| 1.3 | Reactive sulfur species | 24 |
| 1.4 | Reactive halogen species | 4 |
| 1.5 | Other metabolic reactions | 19 |
| 2 | Transports | 129 |
| 3 | Exchanges | 65 |
| 4 | Demands | 1 |
| 5 | Reactions from Recon 3D Reconstruction https://www.vmh.life/ | 62 |

The glutathione redox status of the cancer cells can be computed by experimentally measuring the ratio of concentrations of reduced glutathione (GSH) and oxidized glutathione (GSSG) [35]. In this work, the glutathione redox status was computationally determined by adding the demand reactions for GSH and GSSG and then measuring the flux span ratio of the demand reactions of the GSH and GSSG in each of the models. The glutathione redox status of cells, reduced glutathione(GSH)/oxidized glutathione(GSSG), of RS-CRC was 50% of its value for the CRC metabolic model.

### 3.3 RS module improved the in-silico predictions of CRC model

After integrating the RS module with CRC, FSr were compared across the RS-CRC vs. CRC model and RS-CRC vs. RS-Colon. The details of the number of deregulated reactions are shown in Figure 4. The results of FSr analysis discussed in this section to understand the deregulated metabolic pathways and related reactions in RS-CRC model discussed in this paper are all unique to the CRC and colon models with RS module integrated to them, and they were not highlighted in the analysis of CRC vs. Colon metabolic models. The metabolic pathways enriched on addition of RS module to CRC are shown in Figure 5. Around 50% of the deregulated reactions belongs to these 5 metabolisms. These metabolic reactions play significant roles in the oxidative stress mediated metabolic rerouting in the CRC [36, 37]. The urea cycle reactions had spermine and spermidine as reactants and released hydrogen peroxide as a byproduct, which contributed to the RS pool. An analysis from COBRA Toolbox called as flux enrichment analysis (FEA) [20] is shown in Table 6. It uses hypergeometric 1-sided test and FDR correction for multiple testing to identify the significantly altered metabolic subsystems of GSMMs. Deregulated reactions that are obtained from FSr and sampling analysis for the 2 cases namely RS-CRC vs. CRC and RS-CRC vs. RS-Colon were used in FEA analysis.

**Fig 4.**
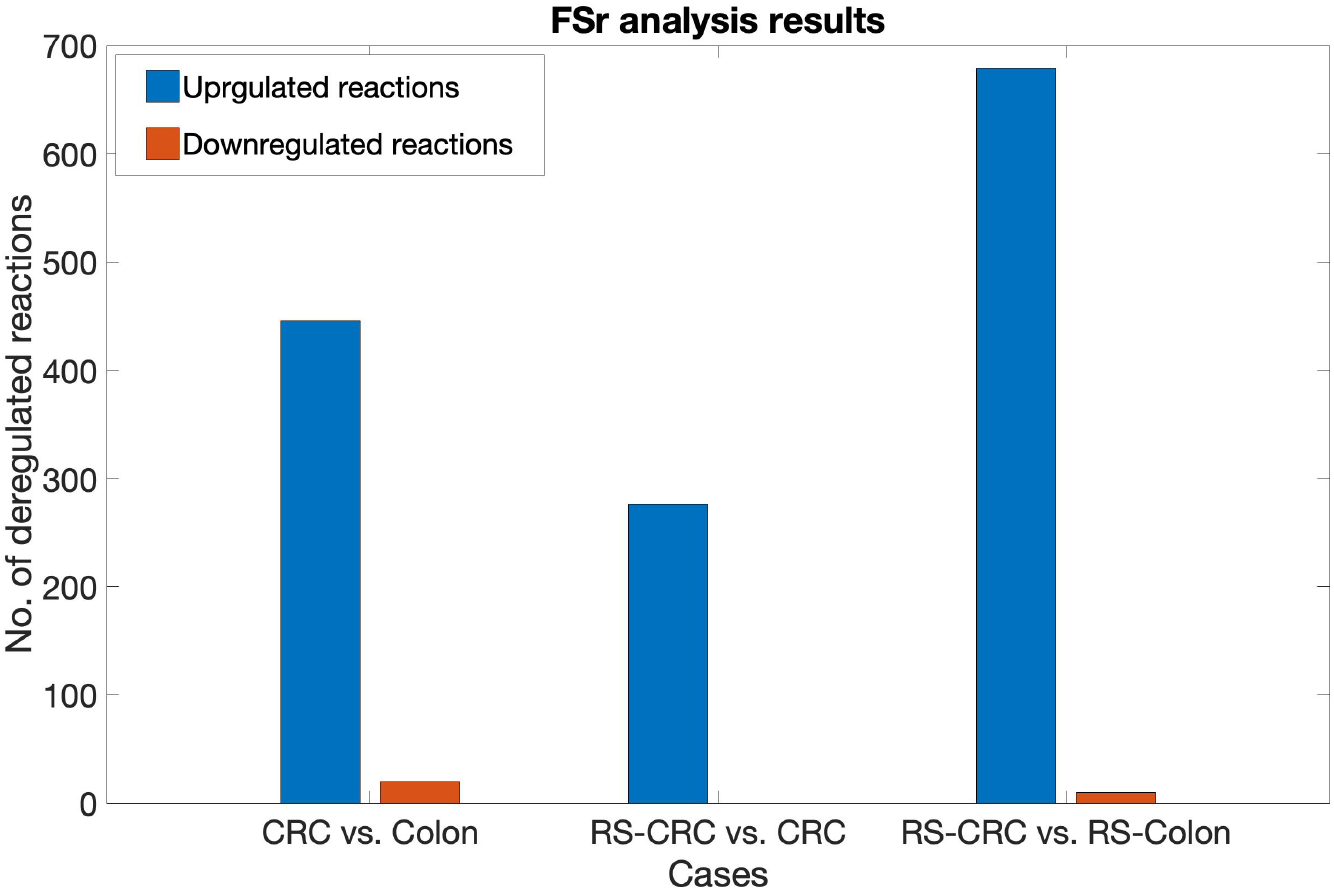
FSr analysis gives the list of deregulated reactions in RS-CRC and CRC models

**Fig 5.**
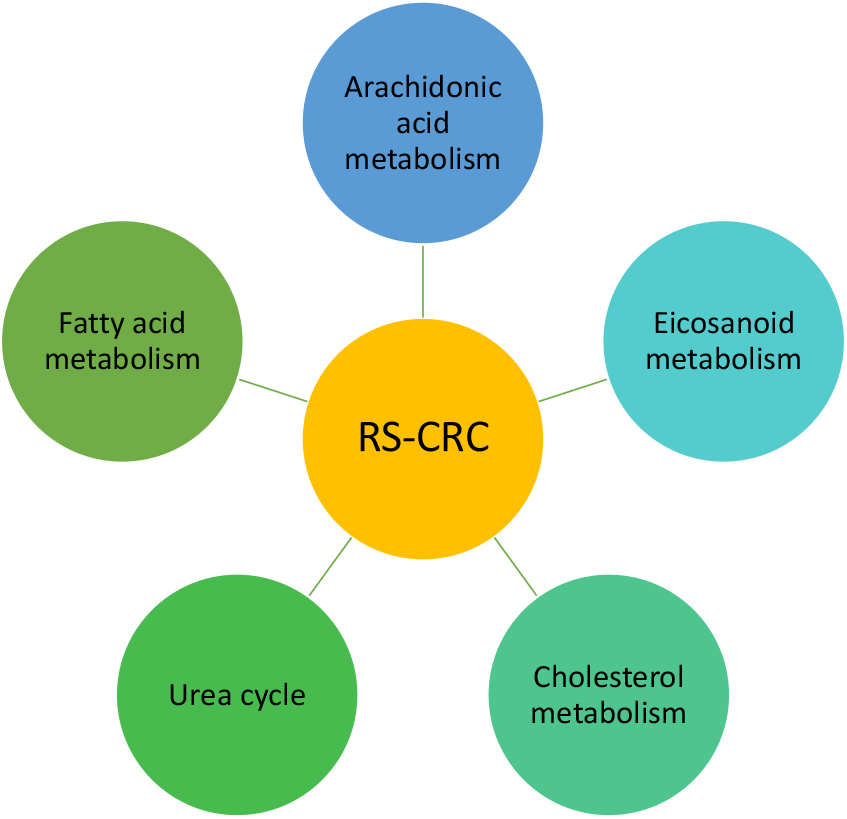
Enriched metabolic pathways that conforms to lipid metabolic reprogramming and redox imbalance in CRC

Numerous enzymes from the important metabolic pathways showed elevated activity (flux in mmol/g-DW/h) in the desired range (given in the methods section) in the presence of RS in the CRC model. The notable ones listed according to the relevant metabolisms are amino acid metabolism (NO synthase, catechol O-methyltransferase); fatty acid metabolism (enoyl coenzyme A hydratase, carnitine O-palmitoyltransferase); steroid metabolism (17-beta-hydroxysteroid dehydrogenase, steryl-sulfatase, steroid 21-hydroxylase, cytochrome P450 17A1); eicosanoid metabolic pathway (prostaglandin endoperoxide synthase 2, prostaglandin I2 synthase); arachidonic acid metabolism (arachidonate 15-lipoxygenase and unspecific monooxygenase). There were no downregulated reactions in CRC after the addition of the RS module.

**Table 2.** Enriched metabolic pathways.

| S.No | RS-CRC vs. CRC | RS-CRC vs. RS-Colon |
| --- | --- | --- |
| 1 | Extracellular transport reactions | Extracellular transport reactions |
| 2 | Fattyacid oxidation | Fattyacid oxidation |
| 3 | Cholesterol metabolism | Cholesterol metabolism |
| 4 | Arachidonic acid metabolism | Arachidonic acid metabolism |
| 5 | Eicosanoid metabolism | Eicosanoid metabolism |
| 6 | Urea cycle | Urea cycle |
| 7 | Steroid metabolism | Bile acid synthesis |
| 8 | Androgen and estrogen synthesis and metabolism | Fattyacid synthesis metabolism |
| 9 | Endoplasmic reticulum transport reactions | Glycerophospholipid metabolism |
| 10 | Tyrosine metabolism | Urea cycle |
| 11 | - | Beta-alanine metabolism |
| 12 | - | Purine synthesis |
| 13 | - | Nucleotide interconversion metabolism |
| 14 | - | RS module reactions |

After comparing the fluxes through CRC and RS-CRC models, subsequent FSr analysis for RS-colon and RS-CRC integrated metabolic models was carried out. From this analysis, some metabolic pathways such as fatty acid oxidation, purine synthesis, fatty acid synthesis, fructose and mannose metabolism, glycosphingolipid and sphingolipid metabolism propanoate metabolism, aminoacids like tyrosine, valine, leucine, and isoleucine metabolism and urea cycle shared a high similarity with the comparison of CRC vs. colon models without RS module integrated with them. The ensuing metabolic pathways were specifically affected in CRC metabolic model in the presence of RS - cholesterol and squalene metabolism (farnesyl-diphosphate farnesyltransferase, lanosterol synthase, squalene epoxidase, 24-dehydrocholesterol reductase, methylsterol monooxygenase, lathosterol oxidase, 7-dehydrocholesterol reductase, etc.); aspartate metabolism (argininosuccinate synthase); linoleate metabolism (linoleate 13S-lipoxygenase, microsomal epoxide hydrolase).

Hypercholesterolemia is a risk factor for colorectal cancer [38] as cholesterol biosynthesis can help in membrane biosynthesis for newly formed cancer cells. Increased levels of oxysterols, which are the oxygenated derivatives of cholesterol is observed in the RS-CRC model when compared against RS-Colon and CRC models. Oxysterols have pro-inflammatory and anti-proliferative roles in colorectal cancers based on the stuidies in CRC cell lines and are known to generate reactive species [39].

Let us next consider the most important pathway, fattyacid metabolism, the following enzymes were found to be dergulated in RS-CRC model, namely carnitine O-palmitoyltransferase, 3-hydroxyacyl coenzyme A dehydrogenase, acetyl coenzyme A C-acyltransferase, enoyl coenzyme A hydratase, acyl coenzyme A oxidase, fatty acyl coenzyme A desaturase, stearoyl coenzyme A 9-desaturase, very-long-chain 3-oxoacyl coenzyme A synthase, etc. Increased fattyacid metabolism observed in RS-CRC plays a significant role in relieving the stress associated with increased lipid accumulation and lipid peroxidation and also in tumour proliferation in colon [40]. The results are shown in the Figure 6

**Fig 6.**
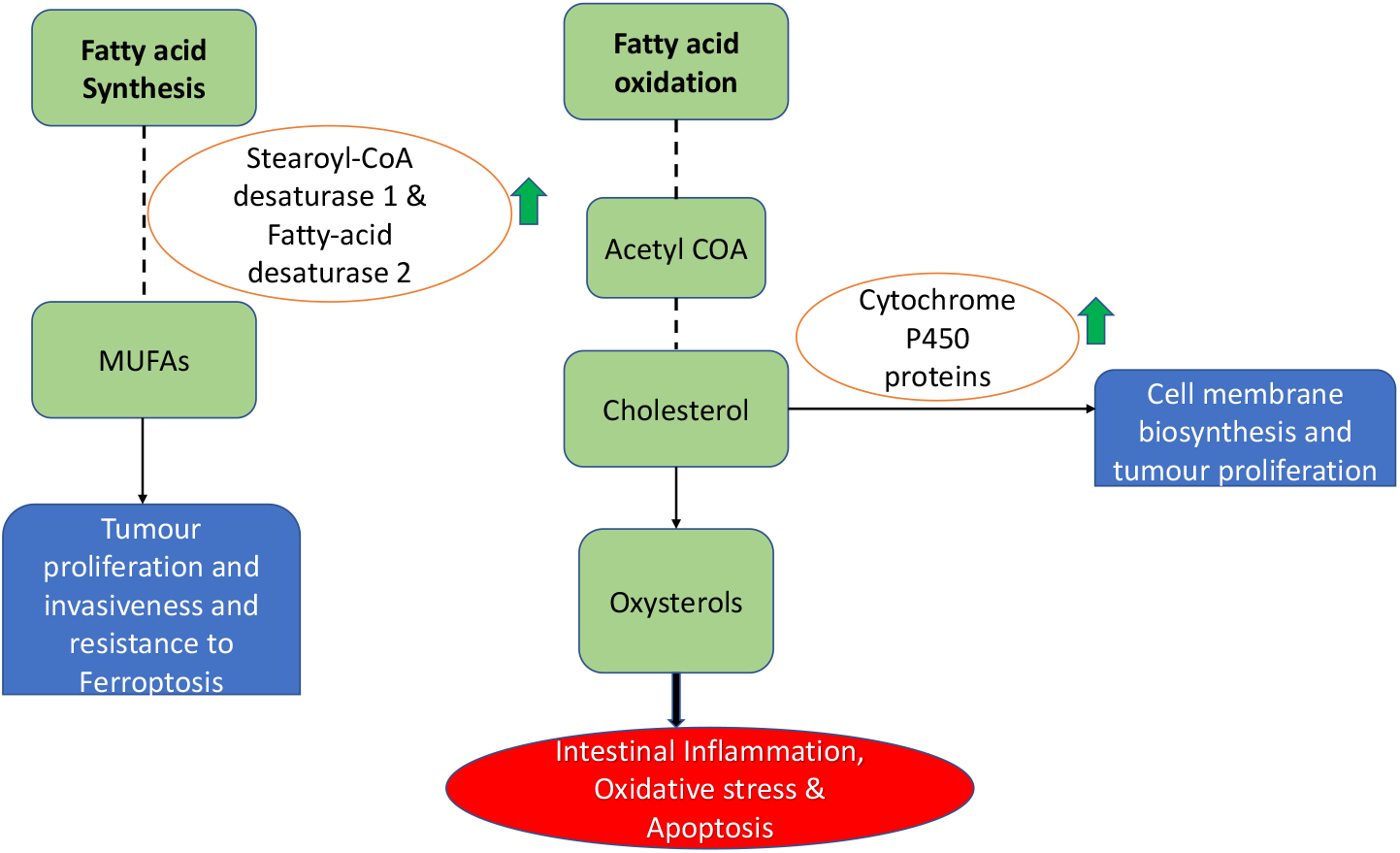
Lipid metabolic reprogramming in CRC

As can be seen in the Table 6, metabolic pathways associated with fatty acid and its derivatives are have undergone significant deregulation in the RS-CRC model. Important enzymes upregulated in the arachidonic acid and eicosanoid metabolism sre shown in Figure 7. Prostaglandins are implicated in the development of CRC and prostaglandin-endoperoxide synthase 2 is overexpressed in 50%-80% of all CRCs which can be observed in RS-CRC model [41]. Leukotriene B_4_ formed from the 5-lipoxygenase pathway is upregulated in the CRC tissue and is found to be overexpressed in the RS-CRC model in comparison to both CRC and RS-colon models. Leukotriene B_4_ overproduction in human colon tissue is implicated in the pathogenesis of inflammatory bowel disease (IBD) and it has a huge role in cancer progression as they are known to activate transcription factors and reactive oxygen species [42].

**Fig 7.**
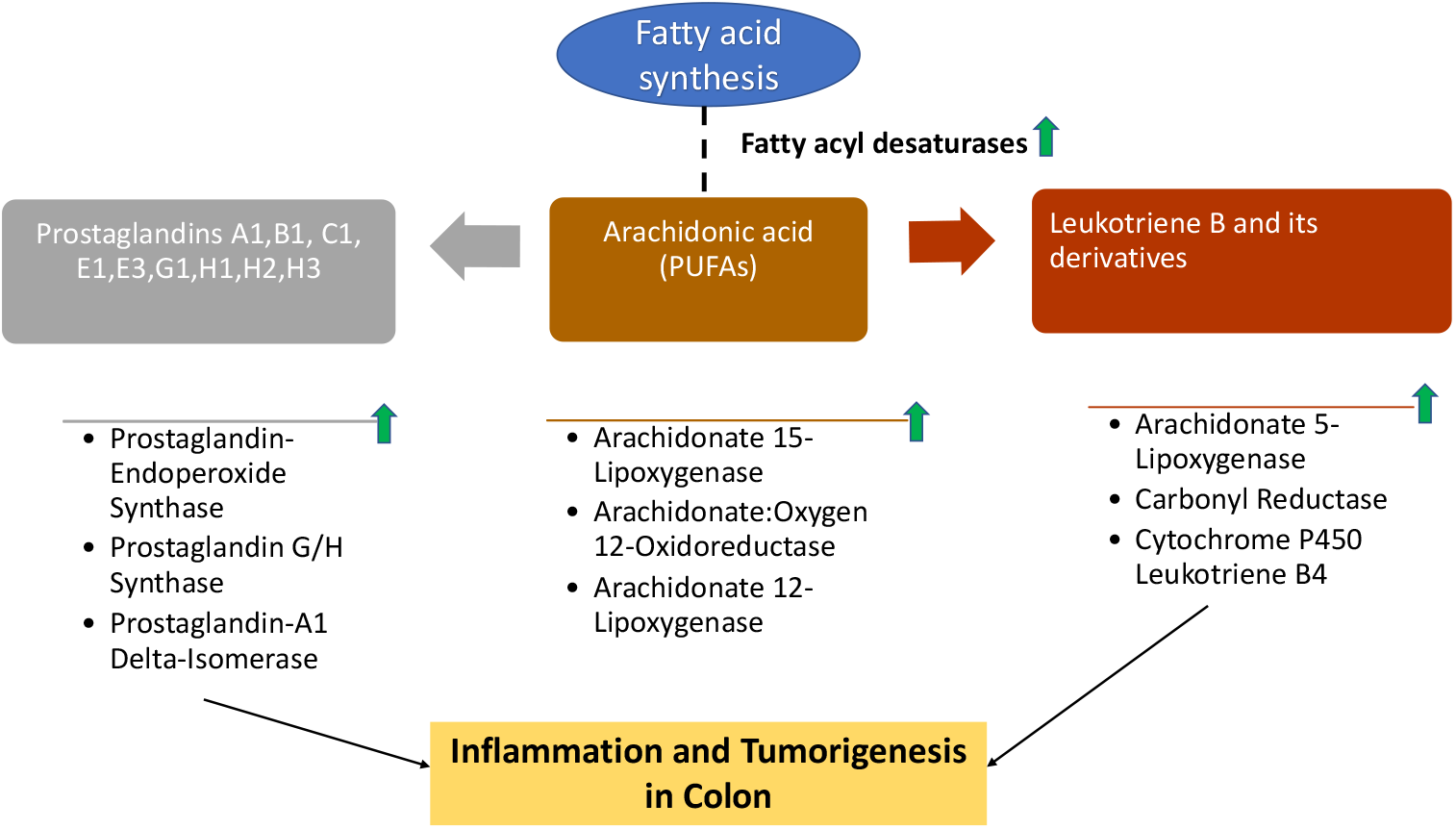
Oxidative stress induces breakdown of ployunsaturated fatty acid especially Arachidonic acid that results in the release of pro-inflammatory molecules like leukotrienes and prostaglandins

#### 3.3.1 Targeting ferroptosis as a weak spot in colorectal cancer

Abnormal lipid metabolism in CRC drives tumor initiation and progression [43]. Ferroptosis is a form of cell death that is prevalent in tumor cells in the presence of iron and lipid peroxides. Reactions associated with the 3 important ferroptosis-related genes namely, farnesyl-diphosphate farnesyltransferase 1 (FDFDT1), arachidonate 12-lipoxygenase (ALOX12) and nitric oxide synthase 2 (NOS2) show increased fluxes in RS-CRC model compared to CRC model. FDFDT1 and ALOX12 associated reactions were upregulated in RS-CRC model compared to RS-Colon. FDFT1 and NOS2 are the protective genes for CRC against ferroptosis and are involved in oxidative metabolism whereas ALOX12 is the risk gene [44]. Metabolism of arachidonic acid by ALOX12 produces 12-hydroxyeicosatetraenoic acid, which has been shown to increase reactive oxygen species in CRC [41]. In spite of the burden from excessive RS species, CRC cells survive by counteracting them with ferroptosis resisting genes, which is captured in our analysis. Even though PUFA synthesis and breakdown happens significantly, MUFA synthesis by FADS2 also occurs in RS-CRC model which is associated with increased resistance to lipid peroxidation and ferroptosis [45]. Thus CRC seems to have protection against ferroptosis mediated cell death, which can be targeted to inhibit CRC development and it is illustarted in Figure 8.

**Fig 8.**
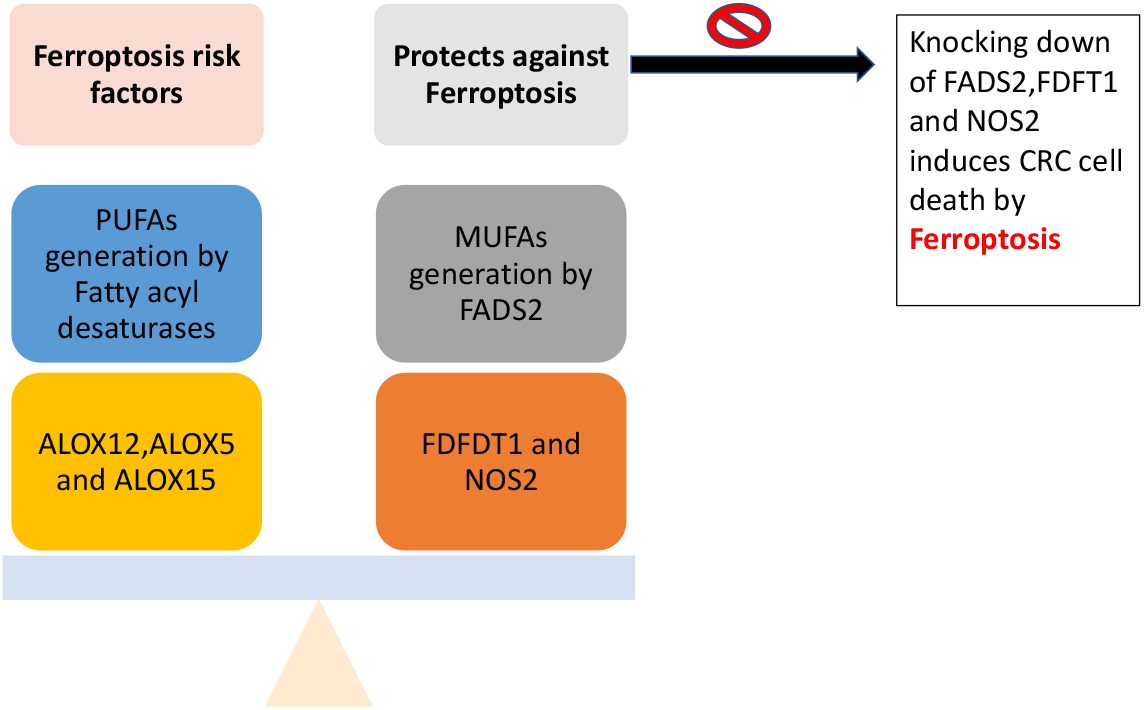
Ferroptosis in CRC

#### 3.3.2 Deciphering metabolic targets in RS integrated CRC by mapping onto HPA druggable proteome

The metabolic (enzymatic) reactions that exhibited pronounced fluxes as observed in the FSr analysis, were further scrutinized for suitable therapeutic drug targets by mapping onto HPA druggable proteome url: https://www.proteinatlas.org/humanproteome/tissue/druggable. A few of these metabolic/enzymatic reactions have been listed as FDA approved drug targets and others as potential drug targets for colorectal cancer by the HPA. Of the numerous reactions that were highlighted in the FSr analysis for RS-Colon vs. RS-CRC models, a few reactions catalyzed by carnitine O-acetyltransferase and carnitine transferase (both belonging to fatty acid metabolism) serve as FDA approved drug targets according to Drugbank https://go.drugbank.com/ for the treatment of colorectal cancer. Moreover, certain reactions from cholesterol metabolism (farnesyl-diphosphate farnesyltransferase), fatty acid oxidation (enoyl-CoA hydratase and 3-hydroxyacyl coenzyme A dehydrogenase) [46] etc., also emerged as potential CRC drug targets. Potential CRC drug targets are those genes that belong to known drug target protein classes of CRC and are not yet approved as targets for FDA approved or experimental drugs in the Drugbank database. Among these targets, farnesyl-diphosphate farnesyltransferase (FDFT1), a potential drug taget for CRC based on evidence from HPA database should be explored further for its efficay in eradicating CRC.

### 3.4 Investigation of in-silico gene targets for devising therapeutics

Single and double gene deletion studies were carried out for the CRC metabolic model, as well as RS integrated CRC model. Compared to the CRC model, the number of lethal, as well as synthetic lethal genes/reactions were considerably reduced in RS-CRC model. Gene and reaction deletion results for CRC and RS-CRC models are shown in the Figure 9. As can be seen from the Figure 9, reduction by more than a third was observed for synthetic lethal reactions deletion, and greater than a 26-fold reduction for synthetic lethal gene deletion. However, some of the lethals as identified in the RS-CRC model were directly or indirectly associated to the reactions involved in the generation of biomass precursors, such as dihydrofolate reductase, phosphatidylserine synthase 1, thymidylate synthase, glycerol-3-phosphate acyltransferase, transport reactions of various nucleic acid biomass precursors, etc. All the lethal reactions of RS-CRC belong to nucleic acid interconversion subsystem and synthetic lethal reactions belong to glycerophospholipid metabolism, nucleotide interconversion, triacylglycerol synthesis, purine catabolism and nuclear and mitochondrial transport reactions.

**Fig 9.**
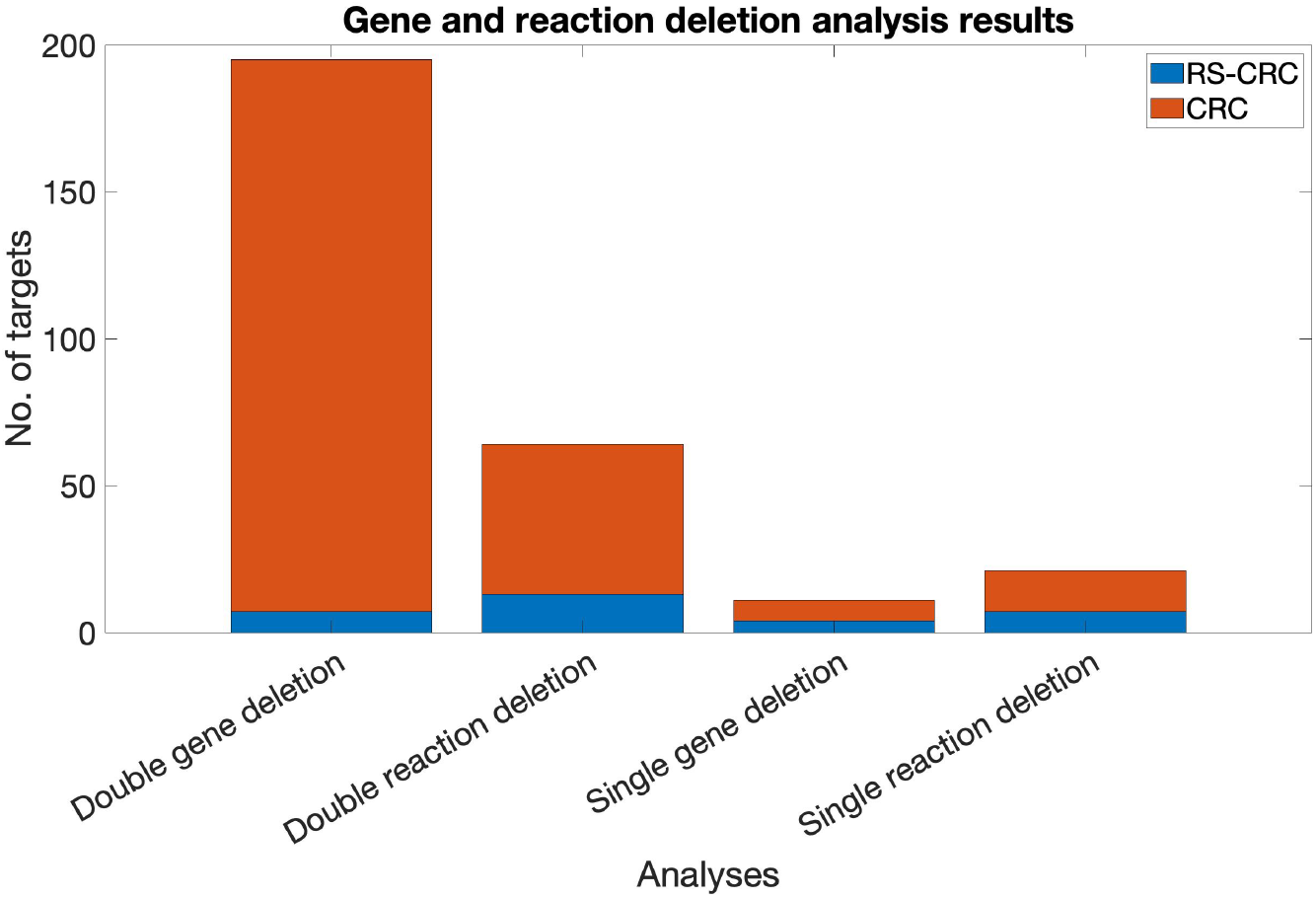
Reactive species reactions increase the resistance of CRC and make them less vulnerable

#### 3.4.1 Validating the predicted in-silico gene targets

The gene targets recognized through FSr and gene deletion analysis were subsequently subjected to validation. To do so, these genes were searched through the DEMETER database [32], which provides information on cancer dependencies and new therapeutic targets (see the methods section). It was observed that most of the genes identified as being significant for CRC cell growth and survival had negative fold change, indicating that they can be further explored as CRC cancer metabolic targets. These genes with their respective log-fold change values are given in Table 7. Novel and potential CRC gene targets obtained from our analysis that were validated against cancer dependency map https://depmap.org/rnai/index and literature were given in the Table 8. Prostaglandin-endoperoxide synthase 2(PTGS2) is a crucial gene in eicosanoid metabolism in colorectal cancers that mediates inflammation and immune suppression [41]. SQLE is a potential therapeutic target for CRC as it promotes CRC by mediating MAPK signaling [47]. Higher expression of LPCAT1 in CRC patients is associated with worse prognosis [48]. Among the genes and gene combinations mentioned in the Table 8, knock down effects of only the first two, PTGS2 and SQLE have been studied in CRC cell lines and animal models [41, 47]. The other proposed targets are found to have an effect on CRC cell line based on the negative log fold values from DEMETER database but are yet to be studied as gene targets in different CRC cell lines.

**Table 3.** Significant in-silico gene targets as identified by FSr/gene deletion analysis and re-confirmed through DEMETER framework.

| Genes | Log-fold change value (DEMETER) | Analysis category |
| --- | --- | --- |
| Carbonyl reductase (CBR1) | -0.180900177 | FSr and Sampling |
| Nitric oxide synthase 1 (NOS1) | -0.17266218 | FSr and Sampling |
| Squalene monooxygenase (SQLE) | -0.10859011 | FSr and Sampling |
| Linoleoyl-CoA desaturase (FADS2) | -0.070496776 | FSr and Sampling |
| Prostaglandin-endoperoxide synthase 2 (PTGS2) | -0.233983909 | FSr and Sampling |
| 1-Acylglycerol-3-phosphate O-acyltransferase (LPCAT1) | -0.092736463 | FSr and Sampling |
| Thymidylate synthetase (TYMS) | -0.415254371 | Lethality analysis |
| Phosphate cytidylyltransferase 1A, choline and phosphate cytidylyltransferase 1B (PCYT1A and PCYT1B), choline | -0.018418591 and -0.139878708 | Synthetic lethality analysis |
| Guanylate kinase 1 (GUK1) and Ribonucleotide reductase M2 B (RRM2B) | -0.340397025 and -0.257455471 | Synthetic lethality analysis |
| Guanylate kinase 1 (GUK1) and ribonucleotide reductase M2 (RRM2) | -0.340397025 and -0.77065238 | Synthetic lethality analysis |
| Guanylate kinase 1 (GUK1) and ribonucleotide reductase M1 (RRM1) | -0.340397025 and -1.005213464 | Synthetic lethality analysis |

**Table 4.** Novel/Potential drug targets uncovered in RS expanded CRC model.

| Genes | Relevance in colorectal cancer | Literature backup |
| --- | --- | --- |
| Prostaglandin-endoperoxide synthase 2 (arachdonic acid and eicosanoid metabolism) | PTGS2 is over-expression is correlated with a lower survival rate among CRC patients | [41] |
| Squalene monooxygenase (cholesterol synthesis) | SQLE stimulates CRC cell proliferation by regulating the cell cycle and its overexpression is associated with poor prognosis in CRC patients in the primary stages | [47] |
| 1-Acylglycerol-3-phosphate O-acyltransferase (fatty acid metabolism) | LPCAT1 stimulates CRC development by enhancing membrane biosynthesis and its a risky prognostic gene in CRC patients | [48,49] |
| Linoleoyl-CoA desaturase (fatty acid synthesis) | Upregulation of FADS2 promotes colon cancer proliferation and tumor growth and yet to be explored as CRC target | [43] |
| Carbonyl reductase (xenobiotics metabolism and antioxidant defense) | Yet to be explored as CRC target and CBR1 overexpression enhances HCC survival by overcoming oxidative stress and cellular stress | [50] |
| Guanylate kinase 1a nd ribonucleotide reductase M1 and M2 (Purine metabolism) | guanylate kinase 1 is yet to be explored as CRC target and is involved in recycling GMP and cyclic GMP and also involved in drug metabolism. Ribonucleotide reductase enzymes are associated with poor survival in CRC | [51,52] |
| Phosphate cytidyltransferase 1A, choline and phosphate cytidyltransferase 1B, choline | Yet to be explored as CRC target and has key roles in phosphatidylcholine biosynthesis and glycerophospholipid biosynthesis | [53] |

## 4 Discussion

### Importance of modelling RS with CRC

RS, especially ROS, are responsible for increased oxidative stress inside a cell, which exert proliferative effects in CRC propagation [54]. Therefore, the primary objective of this study was to model the overall effect and implications of RS on cancer metabolism to develop improved and novel therapeutic strategies. To do so, we developed a scalable RS module using information from databases, and other extensive literature sources, and curated through literature-based information. This RS module can be integrated with any metabolic model to gain insights on the RS effects on the metabolism of those cells. The developed RS module was integrated with tissue (CRC/colon)-specific metabolic models.

The glutathione redox status in terms of GSH/GSSG is an important tumour biomarker of CRC [55] and can be used in non-invasive CRC diagnosis. GSH/GSSG of the CRC model reduced after integrating the RS module to the CRC model due to the increase in the flux activity of GSSG producing reactions. The glutathione redox status of the colon model did not change on addition of RS module, which emphasizes the importance of metabolic reprogramming to compensate for oxidative stress occurring in the CRC metabolic model.

Tumours withstand higher levels of reactive species compared to normal tissues due to their altered metabolism but beyond a point, further elevation in reactive species will disrupt cancer cells. Redox therapy against cancer use this vulnerability to kill cancer kills [56]. In colon cancer, glycolysis, glutaminolysis, fatty acid oxidation (FAO), one-carbon metabolism and the pentose phosphate pathway (PPP) are the 5 major pathways that are upregulated to increase the production of redox cofactors such as NADH, FADH_2_ and antioxidants such as NADPH [56]. However in the RS-CRC model, only fattyacid oxidation is involved in producing reducing equivalents. FAO and fatty acid synthesis are two highly deregulated metabolic subsystems in the RS-CRC model.

### Predictions of FSr analysis on RS-CRC model

Identification of clinical biomarkers will be useful in prediction of diseases at early stages and also will be useful to assess the progress of the CRC over the course of therapy. Thus analysis of serum metabolites becomes essential. We have compared the metabolites in the extracellular[e] compartment of the models and compared it against literature. Conjugated primary (glycochenodeoxycholate) and secondary bile acids (taurodeoxycholate) in the plasma were associated with increased colon cancer risk [57]. Bile acid signalling was known to stimulate cyclogenase 2 enzyme (COX-2) which produces Prostaglandin E which was also highlighted in the RS-CRC model [58]. PGE2 contributes to CRC by inducing fattyacid oxidation associated genes [40] and its level is upregulated in the RS-CRC model compared to RS-Colon model. There are few disagreements between different studies on CRC metabolomics [29, 30] as there is no standard approach of metabolomics studies which result in multiple conclusions. On that note, there are few discripancies with the downregulated metabolites between the literature and our analysis. Nevertheless serum metabolomic profiles are very much essential for diagnosis and treatments. Metabolites associated with reactive species metabolism should also be considered in the future metabolomics experiments as they can give the status of the oxidative stress in the CRC. The altered fluxes in the RS expanded CRC model reactions compared to those of CRC were then analyzed by calculating flux span ratios (FSr). The FSr results emphasized on various metabolic pathways that are deregulated in RS-CRC integrated models. Most of these computational findings on major RS modulated CRC metabolic pathways were confirmed with literature studies. For instance, in the presence of RS, the flux of NO synthase catalyzed reaction showed an increase. Confirming this simulation finding, certain studies have reported increased activity of NO synthase behind CRC proliferation and tumor angiogenesis [59]. Similarly, carnitine O-palmitoyltransferase upregulation is important for tumor promoting effects of adipocytes in colon cancer [60]. Steryl-sulfatase, an essential enzyme required for estrogen activation has also been reported to cause increased CRC proliferation by upregulating a G-protein-coupled estrogen receptor (GPER) pro-proliferative pathway [61], was also affected in presence of RS. Moreover, the RS expanded CRC model also captured the activities of prostaglandins (PGs) and leukotrienes (LTs) synthesizing enzymes - cyclooxygenase (COX) and arachidonate lipoxygenases (ALOXs) (12-lipoxygenase, 15-lipoxygenase and 5-lipoxygenase), which are known to possess heightened activity in CRC [41].

A similar computational analysis was carried out for RS-CRC and RS expanded healthy counterpart of CRC (RS-Colon). The computational findings conformed with literature-based evidence in this case as well. For example, squalene epoxidase participating in cholesterol metabolism has been implicated in promoting CRC proliferation via metabolic and signaling pathways [47]. Moreover, argininosuccinate synthase catalyzed reaction which showed increased flux in RS-CRS model has been associated with CRC pathogenicity by acting as a metabolic regulator [62]. Also, microsomal epoxide hydrolase which deals in metabolizing polycyclic aromatic hydrocarbons and carcinogens found in cigarette smoke and has been linked with development of colorectal adenoma (which progresses to CRC) [63], was also upregulated in RS expanded CRC. Inflammation is an important phenomenon involved in CRC development as inflammatory bowel disease(IBD) is an important risk factor for the development of colon cancer [64]. Prostaglandin endoperoxide synthase 2 is the key enzyme involved in inflammation in CRCs [64]. Non-steroidal anti-inflammatory drugs (NSAIDs) inhibited chemically-induced CRC in rodent models by inhibiting PTGS2 activity [41]. Studies have shown that knockdown of PTGS2 alone or in combination with Arachidonate 5-Lipoxygenase (ALOX5)reduced cellular proliferation in colon cancer [65].

### Predictions of gene deletion analysis on RS-CRC model

Another modeling strategy to uncover potential, as well as existing drug targets in RS-CRC metabolic model is gene deletion analysis (refer methods section). Any deletion of gene(s) or reaction(s) could significantly affect cancer cell biomass. Such single lethal genes/reactions, when targeted, can reduce the flux through biomass reaction to less than 1% of its maximum value, and therefore are crucial for cell survival. This is because such genes/reactions are chiefly responsible for generating biomass precursors like amino acids, membrane lipids, metabolites required for energy maintenance (ATP) and nucleic acids. Likewise, synthetic lethal sets were identified in the cancer model, wherein, pairs of genes/reactions which when deleted together reduce the biomass flux by more than 99% of the original flux value and deleting either of these alone have no such significant effect on biomass. Such pairs of gene/reaction provide a sense of the complex interactions taking place in huge metabolic networks like that of Recon 3D (Supplementary file). Nevertheless, a considerable reduction was observed in the number of gene or reaction targets in the RS-CRC model, when compared to the CRC model alone. For instance, the lethal genes/reactions of CRC model in the absence of RS module belonged to various metabolic subsystems, like NAD metabolism, transport reactions to endoplasmic reticulum and mitochondria, purine catabolism, ubiquinone synthesis and oxidative phosphorylation the related enzymes for which were missing in the gene/reaction deletion analysis of RS-CRC model. This could be attributed to the metabolic re-programming that occurred upon inclusion of RS, which subsequently eliminated the dependency of the CRC cells on these metabolic pathways for their ATP demand. Reactions from the nucleotide intercoversion metabolism, pyrimidine synthesis and glycerophospholipid metabolism are the only three metabolic pathways associated with the synthetic lethals of RS-CRC. Guanylate kinase 1 with Ribonucleotide reductase M1, M2 and M2B and Phosphate Cytidylyltransferase 1A, Choline and Phosphate Cytidylyltransferase 1B, Choline are the potential synthetic lethal targets of RS-CRC model which can be explored for their significance in CRC tissues.

### Ferroptosis is a vulnerability in CRC

The findings from FSr and gene deletion analysis were reconfirmed using the DEMETER framework and suitable literature, thereby ascertaining the credibility of the disease specific RS expanded metabolic model Table 8. The Cancer Dependancy Map is a reliable source of information on large pan-cancer CRISPR-Cas9 gene dependency data sets [66]. The RS-CRC model featured heightened activity of certain enzymes, like farnesyl-diphosphate farnesyltransferase and presqualene-di-diphosphate synthase, encoded by gene FDFT1 which is currently being evaluated as drug target in cancer treatment and is reconfirmed through many sources [67]. Lipid peroxidation induced ferroptosis can increase oxidative stress and iron accumulation and can be targeted in killing colorectal tumour. FDFT1 gene overexpression protects colon cancer cells against ferroptosis and can be a very good drug target to increase sensitivity in ferroptosis resistant cells [68]. From the Figure 8, it can be observed that there exist a fine balance between ferroptosis resisting and ferroptosis promoting mechanisms in the RS-CRC model. Creating an imbalance favouring the ferroptosis promoting mechanisms will help in inducing sensitivity to CRC cells. Knocking down genes such as FDFT1 and FADS2, which are the two central enzymes in RS-CRC that regulate cholesterol metabolism and fatty acid synthesis metabolism can bring down CRC development in patients.

## 5 Conclusion

Constraint based modeling approaches under steady state kinetics are the best bet to understand adaptive antioxidant response against oxidative stress occurring in cancers because they mimic the antioxidant systems of the cells working towards eliminating the RS generated. To summarize, the substantial extent of agreement between the modeling predictions and literature reports has ascertained the credibility of the add-on RS module for providing improved insights into CRC metabolism. Furthermore, when the models were subjected to diverse modeling analysis techniques, the addition of reactive species aided in understanding the disease metabolism, thereby providing directions to identify and establish potential drug targets. Moreover, the RS module can also be integrated with any diseased metabolic models like Alzheimer’s, metabolic conditions like diabetes and inflammatory bowel disorders and other cancer types to assist in studying the relevance of the former in targeting and managing these disorders.

## Supplementary file

**Table 5.**
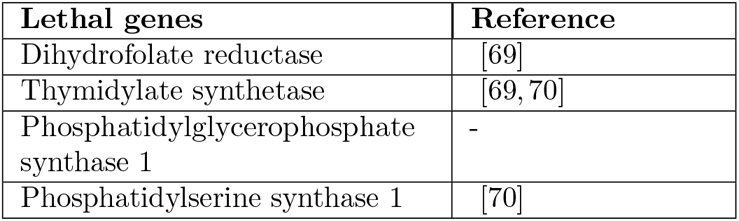
Single gene deletion targets in RS-CRC model.

| **Lethal genes** | **Reference** |
| --- | --- |
| Dihydrofolate reductase | [69] |
| Thymidylate synthetase | [69, 70] |
| Phosphatidylglycerophosphate synthase 1 | - |
| Phosphatidylserine synthase 1 | [70] |

**Table 6.** Double gene deletion targets in RS-CRC model.

| **Synthetic Lethal genes** |  |
| --- | --- |
| Choline/ethanolamine phosphotransferase 1 | Choline phosphotransferase 1 |
| Phosphate cytidyltransferase 1A, choline [53] | Phosphate cytidyltransferase 1B, Choline |
| Deoxythymidylate kinase [71] | SLC19A25 |
| Guanylate kinase 1 | SLC19A25 |
| Guanylate kinase 1 | Ribonucleotide reductase M2 B [51] |
| Guanylate kinase 1 | Ribonucleotide reductase M2 [51] |
| Guanylate kinase 1 | Ribonucleotide reductase M1 [52] |

**Table 7.** Single Reaction deletion targets in the RS-CRC model.

| **Lethal reactions** |
| --- |
| 2-Deoxyadenosine 5-diphosphate:oxidized-thioredoxin 2-oxidoreductase |
| 2-Deoxyguanosine 5-diphosphate:Oxidized-thioredoxin 2-oxidoreductase |
| 2-Deoxycytidine 5-diphosphate:Oxidized-thioredoxin 2-oxidoreductase |
| 2-Deoxyuridine 5-diphosphate:Oxidized-thioredoxin 2-oxidoreductase |
| Ribonucleoside-diphosphate reductase (ADP) |
| Ribonucleoside-diphosphate reductase (GDP) |
| Ribonucleoside-diphosphate reductase (CDP) |
| Ribonucleoside-diphosphate reductase (UDP) |

**Table 8.** Double reaction deletion targets in RS-CRC model.

| Synthetic Lethal reactions |  |
| --- | --- |
| DCTP Diffusion in nucleus | Deoxycytidine nuclear transport |
| Transport of dTTP, Nuclear | Transport of dTDP, Nuclear |
| Phosphatidylglycerol Phosphate phosphatase | Transport of Phosphatidyl glycerol phosphate, Mitochondrial |
| Phosphatidylglycerol phosphate phosphatase | Transport of Phosphatidylglycerol, Endoplasmatic Reticulum |
| Phosphatidyl-CMP: Glycerophosphate phosphatidyltransferase | Phosphatidate cytidyltransferase |
| Phosphatidyl-CMP: Glycerophosphate phosphatidyltransferase | Cytidylate kinase (CMP), Mitochondrial |
| Phosphatidyl-CMP: Glycerophosphate phosphatidyltransferase | Nucleoside-diphosphate kinase (ATP:CDP), Mitochondrial |
| Phosphatidyl-CMP: Glycerophosphate phosphatidyltransferase | CDP-Diacylglycerol-glycerol-3-phosphate 3-phosphatidyltransferase |
| Guanylate kinase (GMP:ATP) | 2-Deoxyadenosine 5-diphosphate:Oxidized-thioredoxin 2-Oxidoreductase |
| Ribonucleoside-diphosphate reductase (ADP) | Deoxyadenosine kinase |
| Ribonucleoside-diphosphate reductase (ADP) | Purine-nucleoside phosphorylase (Deoxyadenosine) |
| Glycerol-3-phosphate acyltransferase | R group phosphotase 3 |
| Lipase | Glycerol-3-phosphate acyltransferase |

## 6 Acknowledgements

The contributions of Sai Sri Teja Thirunahari, Dev Nandan Kumar, Sabyasachi Mishra, Kavan Savla, and Professor Raghunathan Rengaswamy toward the work are acknowledged. Funding from SERB, DST, India, Project number CRG/2020/000119, is gratefully acknowledged.

